# Fungal microbial enrichment method enables fungal metagenomics directly from human clinical samples

**DOI:** 10.64898/2026.07.16.738568

**Authors:** Michael K. Porter, Reid T. Akana, Anna E. Romano, Xinyue (Penny) Pei, Bishoy Kamel, Sajeet F. Haridas, Kurt LaButti, Igor V. Grigoriev, Natalie J. Wu-Woods, Omai Garner, David Underhill, Rustem F. Ismagilov

**Affiliations:** Division of Chemistry & Chemical Engineering, California Institute of Technology 1200 E. California Blvd., Pasadena, CA, 91125 United States; Division of Biology & Biological Engineering, California Institute of Technology 1200 E. California Blvd., Pasadena, CA, 91125 United States; U.S. Department of Energy Joint Genome Institute, Lawrence Berkeley National Laboratory, 1 Cyclotron Road, Berkeley, CA 94720, United States; Department of Plant and Microbial Biology, University of California Berkeley, Berkeley CA 94720, United States; Department of Pathology and Laboratory Medicine, David Geffen School of Medicine at UCLA, 1P-171 Center for Health Sciences, Los Angeles, CA 90095-1732, United States; Department of Biomedical Sciences, Cedars-Sinai Medical Center, 8700 Beverly Blvd., Los Angeles, CA 90048, United States

**Keywords:** Fungal metagenomics, bronchoalveolar lavage, host DNA depletion, microbial enrichment method, metagenome-assembled genomes

## Abstract

Fungi play important roles in health and disease, but current methods such as culture, PCR, and amplicon sequencing cannot provide genome-level characterization directly from clinical samples. Although metagenomic sequencing could overcome these limitations, it remains impractical in clinical samples where fungal DNA is present at low abundance relative to human DNA. Here, we extend a recently described microbial enrichment method (MEM)(1) to fungi (fungal Microbial Enrichment Method; fMEM) and test the method in bronchoalveolar lavage (BAL) samples to demonstrate direct-from-sample fungal metagenomic analysis and metagenome-assembled genome (MAG) recovery. In BAL samples, fMEM depleted human DNA by more than 1000-fold while preserving fungal DNA within 10-fold, enabling shotgun sequencing from samples with fungal biomass as low as 10 pg fungal DNA per 200 µL BAL. fMEM enabled *de novo* recovery of fungal MAGs from three of four sequenced BAL samples, including two near-complete MAGs (>90% BUSCO completeness) and one 82.1% complete MAG, with low BUSCO-estimated contamination (≤1.5%). Fungal MAGs recovered by fMEM also resolved potentially clinically-relevant genes, not fully predictable from taxonomy alone and revealed genomic content absent from currently-available same-species reference genomes. fMEM is compatible with a whole-genome amplification (including long-read sequencing workflows). Long reads from fMEM-processed samples provided high coverage (>10X) over fungal assemblies. fMEM’s compatibility with long-read sequencing enables recovery of genes that would be difficult to assemble with short reads alone. fMEM may enable new insights into the role of human-associated fungi, impacting public health, clinical management, and research into complex diseases with suspected fungal roles.

**Importance:** Fungi influence human health, infectious disease, and the microbiome, but direct genome analysis from clinical samples has remained impractical because fungal DNA is often overwhelmed by human DNA. We developed a fungal microbial enrichment method (fMEM) that enables direct-from-sample fungal metagenomic sequencing and genome recovery from bronchoalveolar lavage samples without requiring culture for genome assembly. fMEM recovers genome-level features not predicted by taxonomy or current same-species reference genomes and is compatible with long-read sequencing workflows that can recover loci missed by short-read sequencing. fMEM opens new opportunities for culture-independent fungal genomics, clinical microbiology, comparative genomics, and mechanistic studies of human-associated fungi.

## Introduction

Fungi play significant roles in human health, infectious disease and the microbiome(2–6). Fungal pathogens cause significant disease burden, ranging from common infections such as candidiasis to deadly infections like aspergillosis. In the human microbiome, some fungi seem to drive disease under certain conditions(7, 8), such as IBD(7, 9), cancer(10–12), and celiac(13, 14). Yet, most human-associated fungi are benign or commensal members of the human microbiome(5, 15–19). Some fungi even seem to contribute positively to health—maintaining gut homeostasis(19), interacting with the immune system(5, 20, 21), or blocking pathogens from invading(19, 22). In some cases, the mycobiome literature is contradictory; a fungal taxon appears protective in some individuals or under certain conditions, but harmful in a different context. Such contradictory findings in the literature emphasize the need for genome-level characterization of fungi directly from clinical samples.

Current methods for fungal detection in clinical samples are either too slow or too confined to predefined targets to provide direct genome-level characterization. The gold standard for fungal diagnostics is culture, which is unacceptably slow (up to 4 weeks)(23) and has poor sensitivity (50%)(24, 25). Culture is also limited to the fungi that can grow outside of the human body, cannot inform on morphologically indistinguishable strains or subpopulations, and may underrepresent genotypic subpopulations that compete poorly in culture(26). PCR is significantly faster and more sensitive than culture-based methods. PCR is routinely used to detect many clinically relevant fungi(27, 28) and some assays also target virulence genes(29, 30). However, PCR still requires a priori knowledge of the target, and primers must be designed and validated for every target in an assay. Amplicon sequencing, which targets loci such as the ITS and 18S genes, yields comprehensive fungal taxonomic information and can sometimes resolve fungi down to species level(31).

However, amplicon sequencing cannot provide strain-level taxonomy, nor can it provide functional information, such as the presence or absence of resistance, virulence, or metabolism genes. Although current methods can provide taxonomic or targeted information, they cannot provide the direct genome-level characterization needed from human-DNA-rich clinical samples.

Direct-from-sample shotgun metagenomics could enable genome-level characterization of fungi, but remains impractical in clinical samples with high human DNA background. Shotgun metagenomic sequencing can overcome these limitations because it analyzes the entire genomic content of a sample. Shotgun sequencing yields millions of DNA sequences (reads) that can be assembled into longer sequences or mapped onto reference genomes, providing strain-level information and the ability to characterize functional capabilities(1). However, performing metagenomic sequencing of fungi directly from clinical samples has been challenging because human DNA greatly exceeds fungal DNA(1). For example, the relative abundances of human DNA can be as high as 95% in saliva(32), 99% in BAL(33), 99.99% in mucosal biopsies(1), and 85% in vaginal swabs(34). In this study, we found the fungal DNA loads in BAL were as low as ∼10 pg/sample, which is 100× lower than the typical input for library preparation for Illumina sequencing. Sequencing more deeply is expensive and still does not overcome the limitations posed by high human DNA loads because the overwhelming majority of the reads will still be human-derived(1). Overcoming this barrier is important for both clinical microbiology and mycobiome research because pathogenic and commensal fungi are often found at low abundance in human-DNA rich mucosal tissues, such as the oral cavity(32), GI tract(35–37), and lungs(33).

Previous direct-sequencing and human DNA-depletion approaches have not enabled fungal genome reconstruction from the most challenging clinically relevant sample types. Although a few previous studies directly shotgun-sequenced human-DNA rich clinical samples to study fungi, they lacked a pre-sequencing fungal-enrichment step, so the resulting sequencing assemblies yielded only taxonomic information(38–41). A wide variety of human-DNA removal (aka “host-depletion”) methods have been developed(32, 42–46), but none have been sufficiently effective to enable fungal MAG construction from sample types containing large amounts of human DNA, such as respiratory samples. The only studies to date that have assembled fungal genomes directly from clinical samples without culture used samples with low human DNA loads, such as stool, where only ∼10% of reads are attributed to human sequences(47).

A technology that depletes human DNA while enriching fungal DNA is therefore needed to enable direct fungal metagenomic sequencing in clinical samples. Genomic characterization of fungi directly from human-DNA rich clinical samples without a culture step for metagenome assembly is critically needed to enable: (1) strain-level taxonomic identification; and (2) identification of potentially clinically-relevant genomic features, such as genes that contribute to virulence, antifungal resistance, biofilm formation, or the ability to colonize. These capabilities would help guide appropriate treatment of infections and greatly improve our understanding of the mechanistic role fungi play in human health(48, 49).

Here, we extend and validate microbial enrichment for fungi by developing the fungal Microbial Enrichment Method (fMEM), a workflow that enables culture-independent fungal metagenomic analysis and MAG recovery from samples with high human DNA background and low fungal biomass. The method leverages our recently developed microbial enrichment method (MEM) that was effective in reducing human DNA by over 1000X in tissue biopsies while preserving bacterial DNA and community composition(1). MEM enabled the first construction of metagenome-assembled genomes of bacteria and archaea directly from samples rich in human DNA, such as human intestinal biopsies. MEM protocol uses selective mechanical lysis followed by proteinase K and benzonase treatment to enrich for bacterial DNA, exploiting the differences in the mechanical properties and size of human and bacterial cells. Fungal cell walls, which are reinforced by chitin, glucans, and glycoproteins, are also more physically robust to mechanical forces than mammalian cells(50–52). Like the MEM protocol, the version of fMEM developed here uses large spherical beads (1.4 mm) to preserve harder and smaller particulate matter, such as fungal cells, whereas standard fungal extraction workflows typically use smaller glass or ceramic beads (0.1–0.5 mm)(53, 54) or garnet beads(55).

To demonstrate the utility of fMEM to enrich fungi in clinical samples, we selected bronchoalveolar lavage (BAL). BAL is an appropriate sample type for testing fMEM because BAL samples are dominated by human DNA (10:1 ratio to fungal cells; up to 1000:5 ratio by genome size; Figure 1B) and known to contain fungal pathogens(33). BAL is also representative of the many clinical sample types that are rich in mucus (e.g., mucosal gut biopsies, vaginal swabs), making it particularly difficult to acquire fungal genomic information. Such samples generally require extra preprocessing, including a longer incubation step with DTT, or saponin to enable consistent sample handling(56).

**Figure 1:**
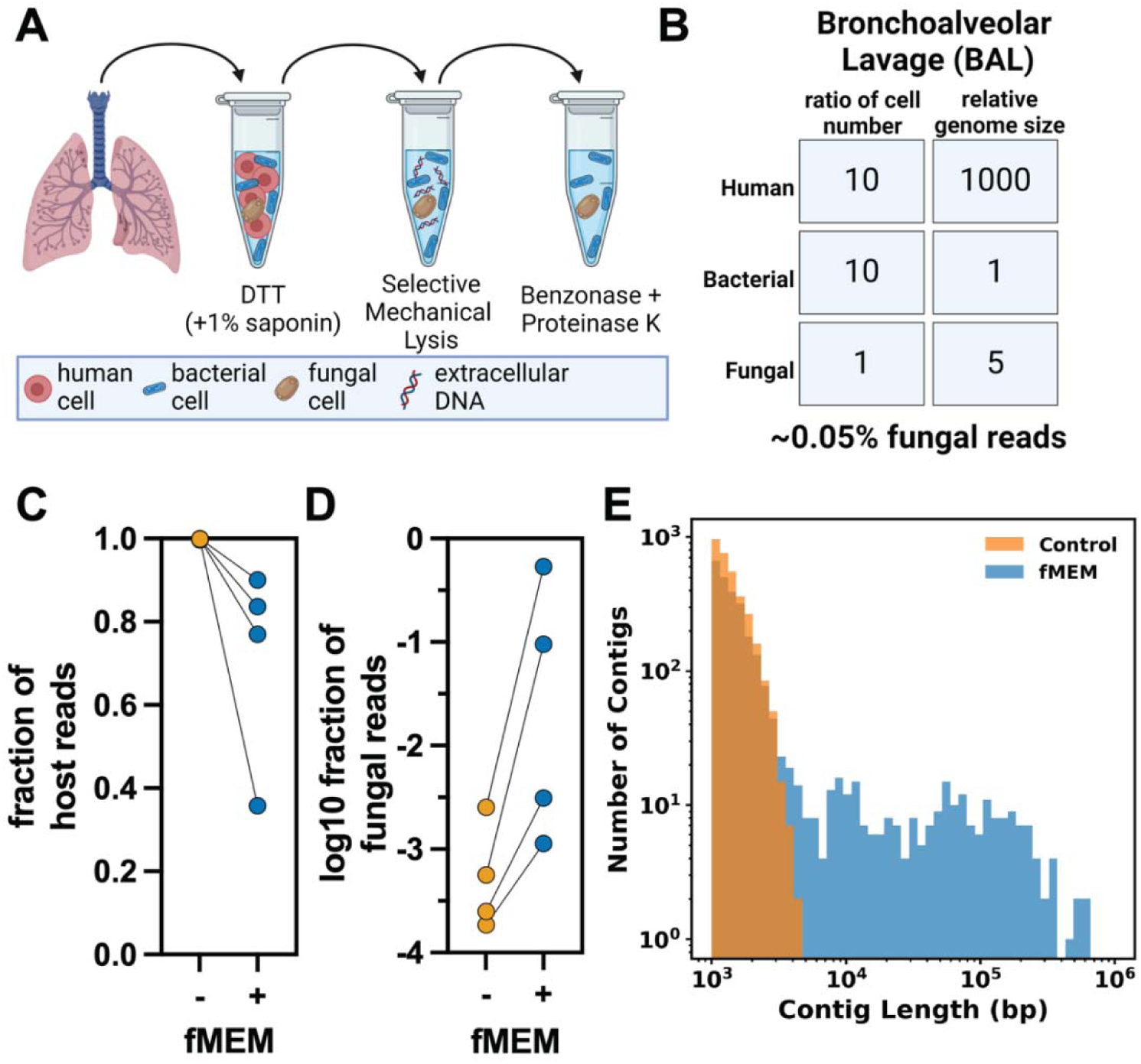
Fungal microbial enrichment method (fMEM) enriches fungal reads and improves fungal contig recovery from bronchoalveolar lavage (BAL) fluid. (A) Schematic of fMEM processing of a BAL sample containing fungi. (B) Estimated cell number and relative genome size ratios of the three main components of BAL (human, bacteria, and fungi), predicting 0.05% fungal reads without enrichment (33). The fraction of reads mapping to human (C) and fungi (D) in BAL with (blue) and without (orange) fMEM. (E) Number of fungal contigs as a function of contig length across four BAL samples with fMEM (blue) and without fMEM (orange).

## Results

### fMEM enriches fungal reads and improves fungal contig recovery from BAL samples

We first asked whether the microbial enrichment method (MEM), previously validated on bacteria(1), could be further optimized to specifically enrich fungi in clinical samples (Figure 1A). We made several modifications to MEM to better enrich for fungi, particularly in challenging mucosal samples (see Methods).

We obtained 22 BAL samples from the UCLA Microbiology Department that were determined to be culture-positive for fungal pathogens. Each sample was processed with and without fMEM, and fungal DNA was quantified by fungal 18S rDNA quantitative PCR (qPCR). Because prior work(57) demonstrated successful Illumina library preparation from as little as 10 pg DNA, we used 10 pg fungal DNA as a threshold for selecting BAL samples for shotgun sequencing. Four of the 22 samples had >10 pg fungal DNA per 200 µL BAL (#011, #148, #160, #170; Figure S1). In these four samples, fMEM reduced human DNA loads by over 1000X, as measured by qPCR (Figure S3), while preserving fungal DNA within 10X (Figure S3).

We next shotgun-sequenced these four BAL samples with and without fMEM processing to determine whether fMEM improved fungal read and contig recovery compared to no enrichment. Both control and fMEM-treated samples were sequenced with Illumina short-read technology to quantify fungal enrichment by fMEM (Table S1). Consistent with the relative enrichment measured by qPCR, fMEM shifted the composition of the sequencing libraries: the fraction of reads mapping to human was 35.8–90.1% (compared with 99.7–99.9% in the controls), and the fraction of reads mapping to fungi was 0.11–53.3% (compared with 0.02–0.25% in controls) (Figure 1C,D).

Additionally, fMEM-processed samples generated fungal contigs that were at least 40 times longer across all four BAL samples than control samples (Figure 1E; Figure S4A–D). Together, these results show that fMEM sufficiently enriches fungal reads and improves fungal contig recovery from BAL to support downstream metagenomic analysis.

### fMEM preserves fungal nucleic acids and community composition

Because fMEM relies on selective mechanical lysis, it could in principle bias recovery toward fungi with more mechanically robust cells. We therefore first asked whether fMEM preserves fungal nucleic acids and does not selectively deplete certain taxa. To test this, we used five cultures of clinically-relevant fungal species and measured both 18S rDNA and 18S rRNA transcripts with and without fMEM using reverse-transcription quantitative PCR (RT-qPCR) (Figure 2A). We included 18S rRNA because RNA is less stable than DNA, so persistence of transcript signal after fMEM provides an indirect proxy for preservation of intracellular fungal nucleic acids. For all species, fungal DNA and RNA losses were approximately 1.32 Cq and 0.834 Cq respectively (Figure 2A).

**Figure 2:**
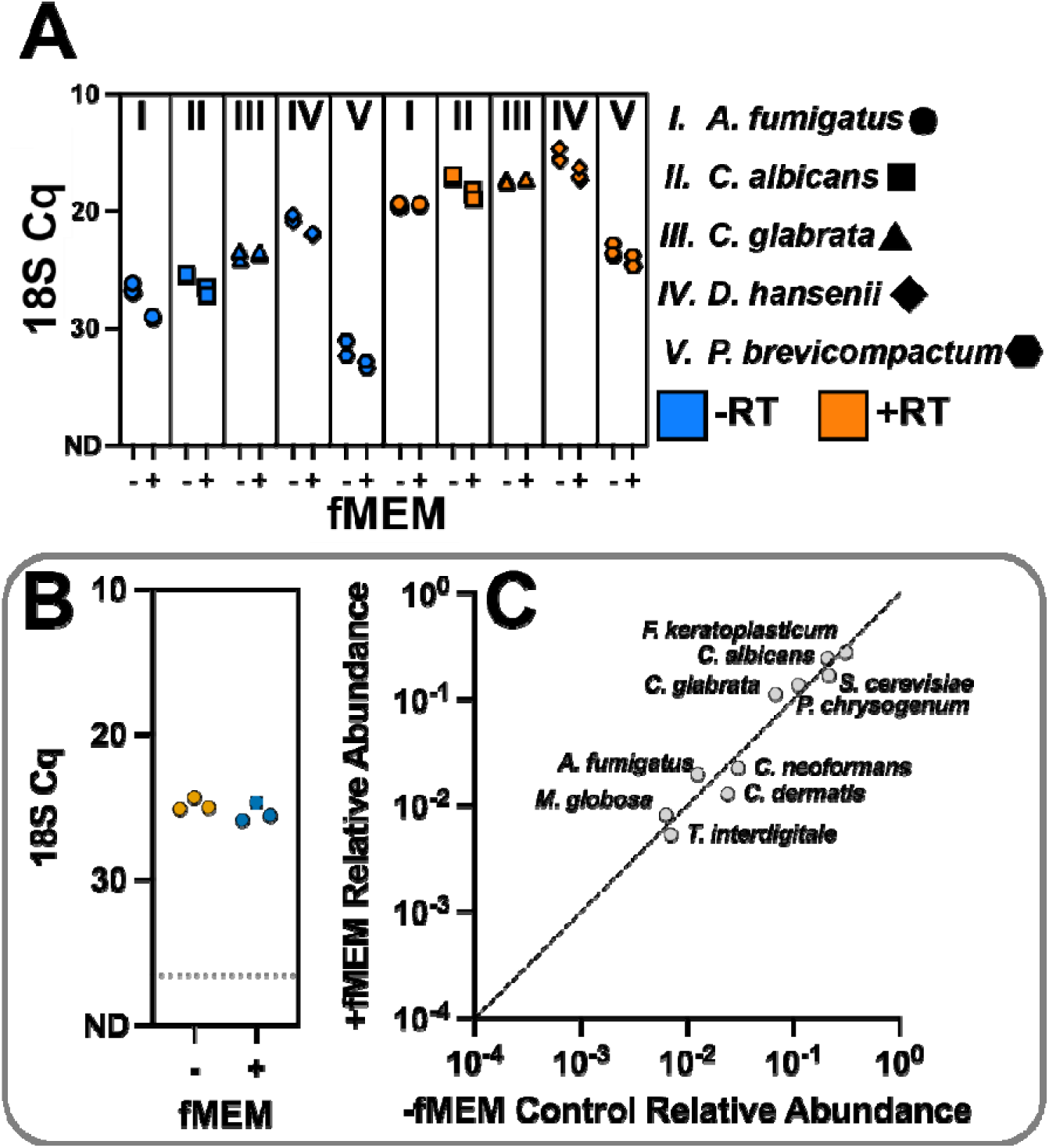
fMEM does not distort the composition of a contrived fungal community standard and preserves fungal DNA and RNA. (A) RT-qPCR Cqs for fungal 18S rRNA transcript (+RT, orange) and fungal 18S rRNA gene (−RT, blue) in fMEM-treated samples (+fMEM) and untreated controls (−fMEM) for (A.I) *Debaryomyces hansenii*, (A.II) *Candida albicans*, (A.III) *Penicillium brevicompactum*, (A.IV) *Candida glabrata*, and (A.V) *Aspergillus fumigatus*. (B) Fungal 18S rRNA gene cycles of quantification (Cq) of the 10-taxa community mix with (blue) and without (orange) fMEM treatment. (C) Relative abundance of each fungal taxon (*Aspergillus fumigatus, Cryptococcus neoformans, Tric ophyton interdigitale, Penicillium chrysogenum, Fusarium keratoplasticum, Candida albicans, Candida glabrata, Malassezia globosa, Saccharomyces cerevisiae, Cutaneotrichosporon dermatis*) after fMEM treatment plotted against its relative abundance before fMEM treatment, measured by 18S rRNA gene sequencing.

We next asked whether fMEM preserves fungal nucleic-acids in a sample with a defined multi-species fungal community standard. We therefore used the 10-taxa ATCC MSA2010 community standard, which includes clinically relevant fungi with a range of physical properties (Figure 2B,C and Figure S2A,B). We first used ITS and 18S qPCR to test whether total fungal nucleic-acids were preserved after fMEM (Figure 2B, Figure S2A), and ITS and 18S amplicon sequencing to test whether fMEM measurably altered the relative composition of the mixed community (Figure 2C, Figure S2B). After fMEM, average losses were 0.56 Cq for fungal 18S and 0.55 Cq for ITS, with high correlation in both assays (18S Pearson r = 0.96; ITS Pearson r = 0.95).

Together, these results show that in contrived fungal samples, fMEM preserved fungal nucleic-acids, with average losses of less than 0.6 Cq. fMEM also did not measurably alter the relative composition of this defined 10-taxa standard, with high concordance between fMEM-treated and control samples by both 18S and ITS amplicon sequencing).

### Enriched fungal reads recruit to paired isolate genomes with high depth of coverage

We next wanted to validate that the fungal reads enriched and sequenced directly from BAL matched the genomes of the cultured fungi present in those samples (i.e., that the recovered genomes were not sequencing artifacts or contamination). To do this, we cultured isolates from the four BAL samples used in Figure 1 for independent validation only; MAGs were assembled directly from BAL and not from cultured isolates.

Three BAL samples had one dominant fungal isolate each: #011 (*Candida lusitaniae*), #148 (*Cryptococcus gattii*), and #160 (*Candida glabrata*). One BAL sample (#170) contained two fungal species (*Candida tropicalis* and *Candida glabrata*).

We sequenced and assembled the genomes from each of these five cultured isolates. All genomes were assembled with high completion (94.8–97.5%) and minimal contamination (0.0–1.5%) (Figure S5). We next recruited the raw fungal reads from the respective BAL samples, with and without fMEM processing, to the paired isolate genomes to confirm that they matched and to compare average depth of coverage (Figure 3). Most reads obtained from the fMEM-processed BAL samples recruited back to the paired isolate genome with moderate to high depth of coverage: 16.91X for #011, 162.28X for #148, 98.15X for #160, and 5.26X for #170 (*Candida glabrata*). In contrast, reads obtained from the control BAL samples recruited to the paired isolate genome with low depth of coverage: 0.03X for #011, 3.95X for #148, 0.31X for #160, and 0.01X for #170 (*Candida glabrata*).

**Figure 3:**
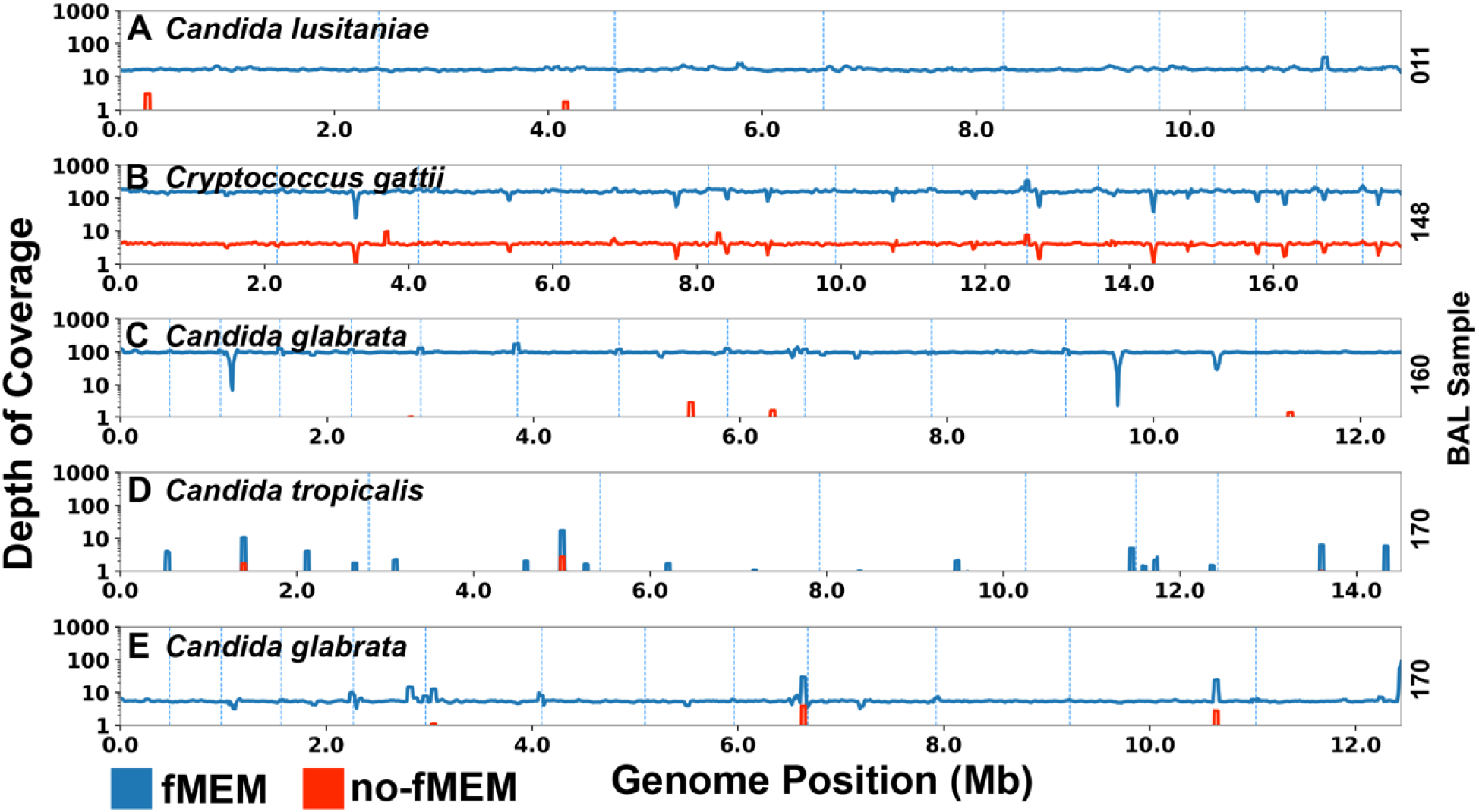
Fungal reads from BAL samples with and without fMEM processing recruit to the paired isolate genomes. Genome-wide depth of coverage after recruitment of reads from fMEM-treated (blue) and untreated (red) BAL samples to the paired isolate genome for (A) #011 (*Candida lusitaniae*), (B) #148 (*Cryptococcus gattii*), (C) #160 (*Candida glabrata*), (D) #170 (*Candida tropicalis*), and (E) #170 (*Candida glabrata*). Quotation marks indicate the species identification assigned by the clinic. #170 contained two fungal isolates.

BAL sample #170 represented a polymicrobial infection case in which one organism remained weakly represented, even after enrichment. For #170 (*Candida tropicalis*), processing without fMEM resulted in an average depth of coverage of 0.00X, whereas fMEM processing increased average depth of coverage to 0.06X. Together, these results show that fungal reads recovered after fMEM correspond to the genomes of the fungi present in the BAL samples and are not sequencing artifacts or contamination.

### fMEM enables direct-from-sample fungal MAG assembly and recovery from BAL samples

We next asked whether fMEM enables recovery of fungal MAGs from BAL samples. We first tested whether fungal MAGs could be assembled *de novo* from fMEM-processed BAL samples. Three of the four BAL samples yielded *de novo* MAGs from 82% to 97% completion (82.1% #011, 97.9% #148, and 92.3% #160, respectively) and low contamination (0.1%, 0.4% and 1.5%, respectively) (Figure 4A-C; dark blue). These *de novo* MAGs had high synteny to the closest-related reference genomes from Mycocosm (#011: MJ12, #148: CA1873) or NCBI (#160: ATCC 2001) (Figure 4A-C; green).

**Figure 4:**
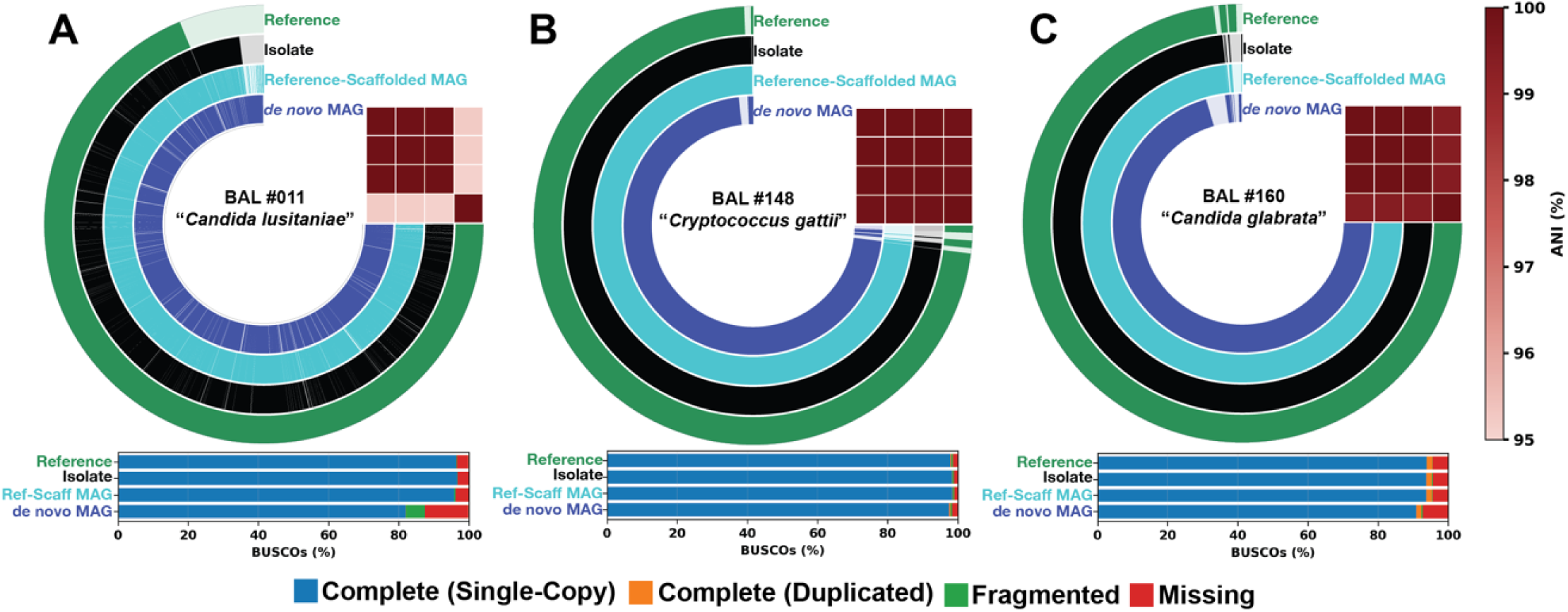
fMEM enables de novo assembly and reference-based scaffolding of fungal MAGs directly from BAL samples. (A–C) *de novo*-assembled MAG (dark blue), reference-scaffolded MAG (light blue), cultured isolate (black), and closest NCBI reference genomes (green) for (A) #011 (*Candida lusitaniae*), (B) #148 (*Cryptococcus gattii*) and (C) #160 (*Candida glabrata*). The de novo MAG assemblies show high concordance with the isolate and reference assemblies. Average nucleotide identity (ANI) is shown as red heatmaps. BUSCO-estimated genome completeness is depicted as a bar graph under pangenome plots for each assembly. Species names indicate the identity of the isolates.

We then generated reference-scaffolded versions of the *de novo* MAGs using MycoCosm or NCBI reference genomes (Figure 4A-C; light blue) and assessed their similarity to the paired isolate genomes (Figure 4A-C; black). The reference-scaffolded MAGs had high Average Nucleotide Identity (ANI) to the isolate genomes (99.98% for #011, #148 and #160). In contrast, the reference-scaffolded MAGs had slightly lower ANI to the reference genomes (95.17%, 99.91% and 99.42% for #011, #148 and #160, respectively), suggesting they are genetically more similar to the cultured isolate from the same sample than to the closest-related reference genome.

We next asked whether fungal MAGs could still be recovered in the BAL samples where *de novo* assembly was not possible. For the one BAL sample where *de novo* MAG assembly was not possible (#170), we used an NCBI reference genome to scaffold fungal contigs from the BAL sample into a MAG (Figure S6). With this reference-based scaffolding approach, a partial MAG for *C. glabrata* from #170 was generated at 56.4% completion with 0.3% contamination. For BAL sample #170, the reference-scaffolded MAG was slightly more similar to the genome of its paired isolate than to the NCBI reference genome (ANI values of 99.87% vs 99.13%) (Figure S6).

Together, these results show that fMEM enables direct-from-sample fungal MAG recovery from BAL samples, including *de novo* MAG assembly in higher-coverage cases and reference-guided recovery of partial MAGs in more challenging cases where *de novo* assembly isn’t possible.

### Direct-from-sample MAGs from fMEM recover orthogroups absent from available same-species reference genomes

Having shown that fMEM enables direct-from-sample fungal MAG recovery, we next asked whether these MAGs reveal genomic features that are incompletely represented in currently available reference genomes of the same species. We compared the #148 MAG and paired isolate assembly with annotated *Cryptococcus gattii* reference genomes from JGI’s MycoCosm database. We focused this analysis on #148 because it yielded a high-quality MAG and paired isolate assembly suitable for orthogroup-level comparison with available *Cryptococcus gattii* references (CA1873, VGII, VGIV, WM276).

In #148, the direct-from-sample MAG and paired isolate recovered expected *Cryptococcus* virulence and antifungal-response genes, such as FKS1(58), FCY1(59), FUR1(60), and RIM101(61). The #148 MAG and paired isolate also carried a small number of variably distributed genes, including a multidrug transporter and phospholipase-family loci, that were present in some, but not all, reference genomes (Figure 5A; orange, green, blue). Further, the #148 MAG and its paired isolate also contained genes absent from all available same-species reference genomes. We found 19 orthogroups that were present in both the MAG and isolate assemblies but absent from the same-species reference genomes (Figure 5A; black/red).

**Figure 5:**
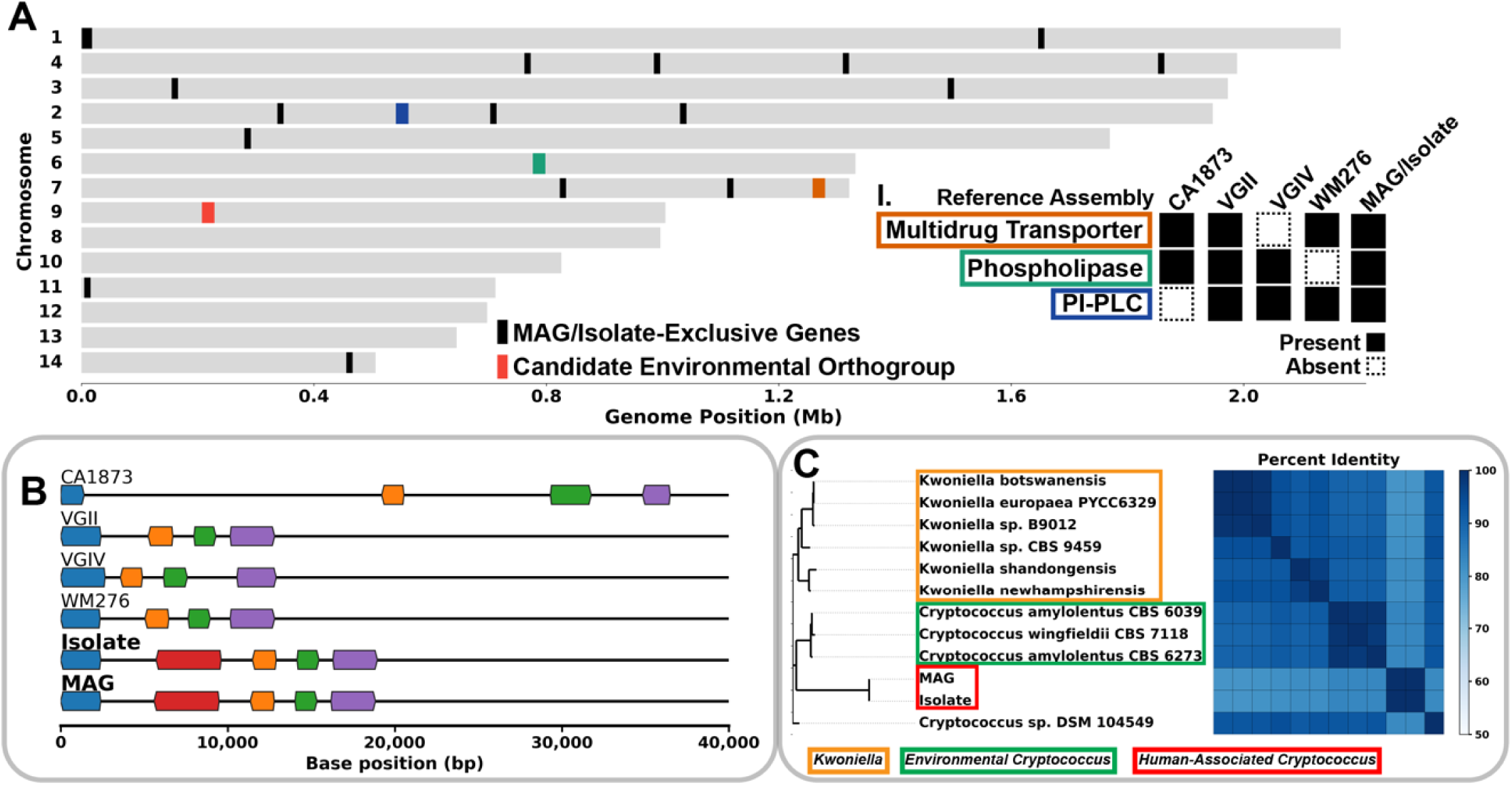
fMEM reveals orthogroups absent from available same-species reference genomes, including an example locus with homologs in environmental *Cryptococcus* lineages. (A) Genome-wide distribution of orthogroups present in the #148 MAG and isolate assemblies but absent from currently-available same-species reference genomes. (I.) Potentially clinically relevant genes identified in the #148 MAG and isolate, including a multidrug transporter (orange), phospholipase (green), and phosphatidylinositol phospholipase C (blue), are also shown, together with their presence (filled squares) or absence (dashed squares) across reference genomes (CA1873, VGII, VGIV, WM276). Red box highlights a MAG/isolate-exclusive locus (OG0006481) selected for further analysis. (B) Gene neighborhood comparison across reference genomes (CA1873, VGII, VGIV, WM276), the isolate genome, and the BAL-derived MAG showing the presence of the OG0006481 (red) locus in the isolate and MAG but not in same-species reference genomes. Conserved flanking orthogroups (blue, orange, green, purple) are shown for context; labels are omitted for clarity. (C) Phylogenetic and sequence similarity analysis (BLASTP) of OG0006481 against NCBI’s nr database shows closest homologs in environmental *Cryptococcus* and *Kwoniella* species.

We selected a representative MAG/isolate-exclusive orthogroup, OG0006481 (Figure 5A; red), for further analysis because its locus was recovered identically in the MAG (protein ID: 58805) and isolate (protein ID: 58824) assemblies. OG0006481 encodes a predicted membrane-associated protein with four transmembrane domains and a glycosyltransferase family 2 (GT2) domain. The presence of conserved surrounding orthogroups (Figure 5B; blue, orange, green, and purple), together with the absence of the OG0006481 locus in the available same-species reference genomes (Figure 5B; red), argues for true presence/absence of the orthogroup rather than misassembly. BLASTP analysis further showed that OG0006481 homologs occur in environment-associated *Cryptococcus* and *Kwoniella* lineages (Figure 5C).

Together, these comparisons indicate that direct-from-sample MAGs can recover conserved genes with potential clinical relevance, resolve variably distributed genes not fully predictable from taxonomy alone, and identify genes present in the sample, but absent from same-species reference genomes.

### fMEM-processed BAL samples support whole-genome amplification and long-read sequencing

We next tested whether fMEM-processed BAL samples could support whole-genome amplification and long-read sequencing, and whether long reads could rescue loci unresolved by short-read assemblies. We selected the three BAL samples (#011, #148, #160) that had already supported *de novo* fungal MAG assembly in the short-read analysis and processed them with primary template-directed amplification (PTA) and PacBio HiFi sequencing (Figures 6A–C) using a library-preparation method we recently developed for direct long-read sequencing of microbial genomes from clinical samples(62). The resulting long-read libraries showed low chimera rates (<2%) and mean read lengths of approximately 4.6 kb (Table S2), and long reads recruited to isolate assemblies with high depth of coverage (>10X) across samples. To test whether long reads enabled recovery of loci that failed to assemble from short reads, we generated hybrid assemblies by re-scaffolding and gap-filling the short-read assemblies with long reads. Although hybrid assemblies showed little global improvement because the short-read assemblies were already highly complete (Table S3), they still rescued specific loci that short reads failed to resolve. For example, in #160, a missing TAF6 BUSCO locus was recovered in the hybrid assembly but was not assembled from the short-read data alone, consistent with long reads spanning a low-complexity region that interrupted the short-read assembly (Figure S7).

**Figure 6:**
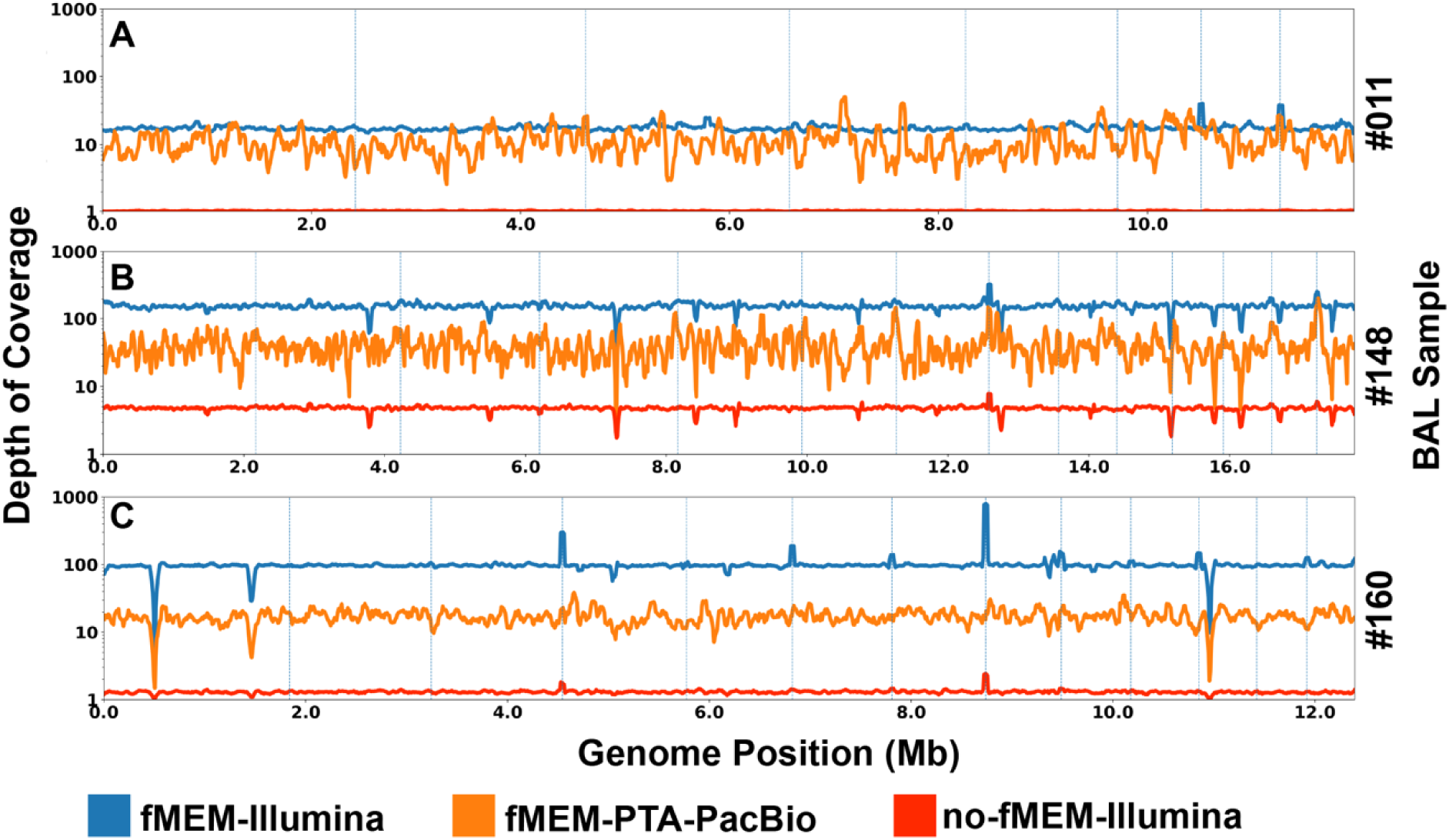
fMEM processing enriches fungal DNA for short- and long-read sequencing. Genome-wide depth of coverage following read recruitment to isolate assemblies for BAL samples with fMEM treatment (blue: Illumina; orange: PTA–PacBio) compared to unenriched controls (red: Illumina). Panels correspond to (A) #011, (B) #148, and (C) #160.

Together, these results show that fMEM is compatible with whole-genome amplification and long-read sequencing, and that the main advantage of long reads in this study was the rescue of specific loci that failed to assemble from short reads.

## Discussion

Here, we developed and demonstrated fMEM, a workflow that enables culture-independent fungal metagenomics from human-DNA-rich BAL samples. fMEM substantially enriched fungal reads and improved contig recovery, overcoming a key barrier to performing direct fungal genome analysis from these samples. fMEM enabled genome-level analysis in low-fungal-biomass clinical samples (∼10 pg) and, to our knowledge, the first direct recovery of near-complete *Candida* and *Cryptococcus* MAGs from BAL.

The direct-from-sample fungal metagenomics enabled by fMEM addresses a key limitation in clinical microbiology and mycobiome research: the ability to go beyond taxonomic profiling to recover potentially clinically relevant gene content – including traits such as antifungal resistance and virulence that are not always predictable from species assignment. For example, in BAL sample #148, the direct-from-sample MAG recovered Cryptococcus genes with potential relevance to virulence and antifungal response. This MAG additionally revealed genes in the patient’s strain that were either sporadically present or completely absent from same-species reference genomes on JGI’s MycoCosm database, including 19 orthogroups absent from all available same-species references. The advantage of direct-from-sample sequencing is especially important for fungi that are difficult or impossible to culture, such as *Malassezia* or *Pneumocystis*.

fMEM was also compatible with whole-genome amplification and long-read sequencing, enabling recovery of difficult loci that were not resolved by short-read amplification/assembly. In this study, fMEM combined with PTA and PacBio sequencing recovered specific hard-to-assemble regions, showing that the main value of long reads was recovery of otherwise missing regions rather than broad improvement of already strong short-read assemblies.

Our study has three main limitations. First, fMEM does not deplete bacterial DNA, which can comprise up to 99% of the biomass in certain clinical samples, such as stool. For some applications, this is a benefit of the method because it enables simultaneous metagenomic analysis of both bacteria and fungi. However, for applications focused on very low-abundance fungi, fMEM could be combined with bacterial–fungal cell-separation approaches(63) to reduce bacterial DNA background. Second, fMEM has been validated only on yeasts and not on hyphal morphotypes, which may be less robust to the mechanical lysis steps and require further optimization. Although most clinically relevant fungi are found as yeasts(64), some applications would benefit from the ability to capture additional fungal morphologies(65). Third, only 4 of 22 BAL samples contained sufficient fungal DNA to proceed to shotgun sequencing, and one fMEM-processed sample did not support *de novo* MAG assembly. Notably, when *de novo* assembly was not possible because of low fungal biomass and high bacterial load, partial fungal MAG assembly could still be achieved via fMEM with reference genomes.

These limitations define several next steps for extending fMEM to broader clinical applications. Here, fMEM was demonstrated on BAL, an important sample type for diagnosing and studying fungal respiratory infections, such as aspergillosis(66). fMEM should be validated in additional human-DNA-dominated sample types, such as vaginal swabs, to enable study of diseases in which fungi play established or suspected pathogenic roles, including vulvovaginal candidiasis. Because this study only validated fMEM on yeasts, future work should also determine how well the method preserves and enriches fungi with diverse morphologies and cell-wall architectures, including septate and aseptate hyphal forms. Additional studies should also define the fungal-load and read-depth thresholds required for *de novo* fungal MAG recovery and determine when long-read sequencing provides enough benefit to justify the added workflow.

Once validated across additional sample types, fMEM will extend direct fungal genomics to clinical settings where culture is slow, biased, or infeasible. More broadly, the ability to recover patient-associated fungal genomes from human-DNA-rich specimens may support clinical microbiology, fungal pathogen surveillance, expansion of reference genomes, and mechanistic studies of how fungi contribute to human disease.

## Materials and Methods

### Human bronchoalveolar lavage (BAL) samples

Human BAL samples were acquired from immunocompromised adults at the University of California, Los Angeles, under UCLA IRB protocol #19-001098. Samples were de-identified and sent to Caltech to be analyzed under California Institute of Technology Institutional Review Board (IRB) protocol #19-0915 (exempt). Before undergoing fMEM, BAL samples underwent a DTT pretreatment (10mM DTT in autoclaved 0.9% NaCl), vortexed, and incubated for 30 min at room temperature before undergoing fMEM. BAL samples that were visibly mucous and opaque underwent DTT (10mM DTT in autoclaved 0.9% NaCl) and 1% saponin treatment at 37C for 30 min.

For cultured isolates, 10 mL of BAL samples were vortexed and centrifuged at 1500 xg for 10 minutes. The supernatant was removed and vortexed for 15-20 seconds before 30-60 uL was added to inhibitory mold agar and sabhi agar. The plates were streaked for isolation and incubated at 30C.

### Cultured single species fungal samples

Single species fungal cultures were prepared at Cedars-Sinai Medical Center in Los Angeles, CA. Culturing conditions for each strain were as follows: *Aspergillus fumigatus* isolate AF293 and *Penicillium brevicompactum* ATCC 9056 were grown on potato dextrose agar (ATCC Medium 337) plates at room temperature. *Candida albicans* ATCC 90028, *Candida glabrata* ATCC 2001, and *Debaryomyces hansenii* ATCC 36239 were grown on yeast extract peptone dextrose (YPD) plates (ATCC media 1245) at 30 °C.

### Fungal Microbial Enrichment Method (fMEM) protocol

Samples that underwent fMEM treatment were processed as previous described(1) with minor modifications to enrich fungi and reduce mucus viscosity. Briefly, BAL samples were thawed on ice and added into 2-ml 1.3-mm ceramic bead beating tubes (Lysing Matrix D from MP Biomedical, catalog no. 116913050-CF) and supplemented with saline (0.9% NaCl, autoclaved) so that the final volume in the bead beating tube was 400 µL. Samples were homogenized using FastPrep-24 (MP Biomedical catalog no. 116004500) for 30 s at 4.5 m/s. A total of 183 µL of homogenized sample was removed and placed into a clean microcentrifuge tube containing 10 µL of buffer (100 mM Tris + 40 mM MgCl2, pH 8.0 and 0.22 um sterile filtered), 5 µL of Proteinase K (NEB catalog no. P8107S), and 2 µL of Benzonase Nuclease HC (EMD Millipore catalog no. 71205). Sample tubes were placed on a dry block incubator for 15 min at 37C while shaking at 600 rpm. Following incubation, samples were centrifuged at 10,000g for 1 min and the supernatant was discarded. BAL samples that were visibly mucous and opaque underwent an additional enzyme wash. Pellets were resuspended in either 750 µL Zymo DNA/RNA Shield (catalog no. R1100-50) for contrived samples or BAL samples and stored on ice until nucleic-acid extraction.

### MAG assembly and Pangenome Analysis

Adapter sequences and low quality/homopolymer sequences were trimmed from reads using fastp(67). Host reads were then filtered using a masked version of the GRCh38(68) human genome with BBMask(69) and BBDuk. Reads were assembled using metaSpades(70). Fungal contigs were identified using Kraken2 and the “PlusPF” database(71) with default confidence thresholds. Fungal contigs were then extracted and used to filter reads. For reference-guided MAG recovery, the closest available fungal reference genome from JGI MycoCosm or NCBI RefSeq was used to identify fungal contigs in the metagenomic assembly, and these contigs were then scaffolded against the reference using RagTag(72).

Contigs were then reassembled with filtered reads using metaSpades. Completeness and contamination of assemblies were assessed using BUSCO with the “fungi_odb10” database(73). Assembly quality was also assessed using genome-wide read recruitment depth, and comparing ANI and synteny to paired isolate and reference genomes. MAG assemblies were additionally screened for non-target contamination using NCBI’s FCS-GX tool(74) and flagged contaminant contigs/sequences were removed before downstream analyses. Depth of coverage was determined for each base in an assembly by recruiting reads back onto assemblies using bwa(75) and depth was calculated with samtools depth(76). Average depth of coverage was calculated by averaging over the depth of coverage of each base in a genome. ANI was calculated across closely related reference genomes obtained from NCBIs RefSeq(77) or JGI’s Mycocosm(78) databases using skani(79) and synteny was visualized using anvi’o(80). For anvi’o pangenome analysis, funannotate2(81) was used to annotate all assemblies. Phylogenomic trees were generated using Orthofinder(82) and iqtree(83) and plotsr(84).

### Hybrid MAG assembly

Long reads were first trimmed with fastplong(67) and host reads were removed with BBDuk(69) using the masked GRCh38 reference described above. Fungal-associated long reads were then identified by mapping long reads against the corresponding fungal short read assemblies. Fungi-associated long reads were assembled with hifi_asm(85). The resulting long-read assemblies were used to rescaffold and gap fill the corresponding short-read assemblies with RagTag(72), generating final hybrid assemblies. Quality of assemblies was assessed with BUSCO(73).

### Gene calling and functional annotation

For orthogroup comparisons, we restricted the reference set to same-species genomes from JGI MycoCosm so that all genomes were annotated with the same JGI annotation pipeline(78, 86). Briefly, RepeatMasker(87) was used in combination with the RepBase library(88) and repeats larger than 150 bp generated by RepeatScout(89) to mask the assemblies. Masked assemblies were staged into the gene prediction pipeline. *Ab initio* gene prediction was used using Fgenesh, Fgenesh+(90) and GeneWise(91) which use seeds from protein to genome alignments. Best models at each locus were selected using the strength of homology support. Functional annotation of the proteins included, using eggNOG-Mapper(92) for EC number assignment, InterproScan(93), for protein domains, TCDB(94) for transporter prediction, TMHMM(95) for transmembrane domains and SignalP(96, 97) for signal peptide prediction, in addition comparisons to NCBI NR, swissprot, KOG(98), KEGG(99) were incorporated. The information from InterPro and Swissprot were used to confidently assign gene ontology terms.

## Acknowledgments

We thank SPEC for access to sequencing instrumentation and Jason Stajich for advice on the manuscript. We also thank Natasha Shelby for contributions to editing this manuscript.

## Funding

This work is supported by an NSF Graduate Research Fellowship (to NWW), a Caltech Beckman Collaborative Pilot Award (to RFI) and a Caltech Resnick Sustainability Institute Impact Grant (to RFI). The work conducted by the U.S. Department of Energy Joint Genome Institute (https://ror.org/04xm1d337), a DOE Office of Science User Facility, is supported by the Office of Science of the U.S. Department of Energy operated under Contract No. DE-AC02-05CH11231 (to IVG). Some of the non-human in vitro and computational development work was supported in part by DARPA contract HR0011263E024 (to RFI); any opinions, findings, conclusions, or recommendations expressed in this material are those of the author(s) and do not necessarily reflect the views of the Defense Advanced Research Projects Agency (DARPA). This manuscript is approved for public release, distribution unlimited.

## Author Contributions

**MKP:** Conceptualization, Methodology, Investigation, Formal analysis, Visualization, Writing – original draft.

**RTA:** Conceptualization, Investigation, Formal analysis, Software, Visualization, Data curation, Writing – original draft.

**AER:** Conceptualization, Methodology, Investigation, Resources, Supervision, Writing – review & editing.

**XPP:** Investigation.

**BK:** Formal analysis, Software, Data curation, Visualization, Supervision.

**SFH:** Formal analysis, Data curation.

**KL:** Supervision, Writing – review & editing.

**IVG:** Formal analysis, Supervision, Writing – review & editing.

**NWW:** Conceptualization, Investigation.

**OG:** Conceptualization, Resources, Supervision.

**DU:** Methodology, Resources, Validation, Supervision.

**RFI:** Conceptualization, Methodology, Supervision, Project administration, Funding acquisition, Writing – review & editing.

### Competing Interest Statement

The technology described in this publication is the subject of a patent application filed by Caltech.

## Data, Materials, and Software Availability

Human-filtered raw sequencing data are available in the NCBI Sequence Read Archive (SRA) under BioProject PRJNA1460817. Genome assemblies and annotations have been deposited in the JGI MycoCosm database under Clavispora lusitaniae MP2BAL, Clavispora lusitaniae MP11, Cryptococcus decagattii MP4BAL, Cryptococcus decagattii MP12, Nakaseomyces glabratus MP6BAL, and Nakaseomyces glabratus MP13. All other data will be made available at CaltechDATA: https://doi.org/10.22002/fevd5-av916

## Notes

### Summary of Updates

Addition of distribution statement required by a funder.

