## Supplementary Information for "Fungal microbial enrichment method enables fungal metagenomics directly from human clinical samples"

**This PDF file includes:**

Supporting text
Figures S1 to S7
Tables S1 to S3
Author Contact Information and ORCiDs
Detailed Author Contribution Statements
Supplementary References

### Supporting Information Text

*Nucleic acid extraction of BAL, cultured clinical isolates, and contrived samples*

Nucleic acids from BAL specimens, contrived fungal communities (ATCC MSA2010), and cultured strains were isolated following Zymo's ZymoBIOMICS MagBead DNA/RNA kit (catalog no. R2135) using the KingFisher Flex 96 instrument (Thermo Fisher Scientific). Samples were first homogenized in a 2-mL tube containing 1.4-mm ceramic beads, 0.1-mm silica spheres, and one 4-mm glass bead (Lysing Matrix E from MP Biomedical, catalog no. 116914050-CF) for 1 min at 6 m/s for three cycles using FastPrep-24 (MP Biomedical catalog no. 116004500), incubating the samples on ice for 5 min between each cycle.

*Quantification of host DNA*

Human host DNA in extracted samples was characterized by quantitative PCR (qPCR) using the Zymo Femto Human DNA Quantification Kit (catalog no. E2005) according to the manufacturer's protocol.

*Quantification of fungal rRNA and DNA*

Fungal rRNA transcripts and DNA in extracted samples were characterized by qPCR. 18S rRNA was first reverse-transcribed using a High-Capacity cDNA Reverse Transcription Kit (Thermo Fisher catalog no. 4368814). Following reverse transcription, 18S cDNA was amplified based on primers adapted from (1). Thermocycling was performed on a CFX96 Real-time PCR system (Bio-Rad Laboratories) using the following protocol: 94 °C for 5 min, 40 cycles of 94 °C for 30 s, 58.3 °C for 50 s, and 72 °C for 1 min. The concentrations of the components in the qPCR mix used in this study were as follows: 1x AccuStart II PCR SuperMix (Quantabio catalog no. 95137-04K), 1x EvaGreen Dye (Biotium catalog no. #31000), 500 nM forward primer, and 500 nM reverse primer for a total reaction volume of 10 µL.

A calibration curve relating fungal 18S assay Cq values to estimated fungal DNA mass was generated using a dilution series of the ATCC MSA1010 10-taxa whole-cell fungal standard:

$$18S\,\frac{ng}{rxn} = 2^{-0.8748Cq+12.175}$$

The ITS DNA was amplified based on the following primers from literature(2): (Forward: 5'- TCCGTAGGTGAACCTGCGG-3' and Reverse: 5'-GCTGCGTTCTTCATCGATGC-3', Integrated DNA Technologies). Thermocycling was performed on a CFX96 Real-time PCR system (Bio-Rad Laboratories) using the following protocol: 94 °C for 5 min, 40 cycles of 94 °C for 30 s, 61 °C for 50 s, and 72 °C for 1 min. The concentrations of the components in the qPCR mix used in this study are as follows: 1x AccuStart II PCR SuperMix (Quantabio catalog no. 95137-04K), 1x EvaGreen Dye (Biotium catalog no. #31000), 500nM forward primer, 500 nM reverse primer, and 1 µL of template for a total reaction volume of 10 µL.

*Fungal 18S and ITS gene amplicon sequencing*

Extracted DNA was amplified and sequenced using 18S and ITS primers described above with Illumina barcode adapter regions (Integrated DNA Technologies). The concentrations of the components in the qPCR mix used for both primers are as follows: 1x KAPA HiFi HotStart ReadyMix (Roche catalog no. 09420398001), 1x EvaGreen Dye (Biotium catalog no. #31000), 500 nM forward primer, 500 nM reverse primer, and 2.5 µL of template for a total reaction volume of 25uL. Amplification was monitored on a CFX96 Real-time PCR system (Bio-Rad Laboratories) and samples were removed once fluorescence measurements reached late exponential phase. The following amplification protocol was used for ITS: 94 °C for 3 min, up to 40 cycles of 95 °C for 30 s, 61.4 °C for 30 s, and 72 °C for 2 min. For 18S: 94 °C for 5 min, up to 40 cycles of 94 °C for 30 s, 62.5C for 50s, and 72 °C for 1min. Illumina barcode ligation was performed after initial amplification using the following protocol: 95 °C for 3min, 8 cycles of 95 °C for 3 min, 55 °C for 30 s, 72 °C for 30 s, followed by 72 °C for 5 min. The concentrations of the components in the reaction mix used are as follows: 1x KAPA HiFi HotStart ReadyMix (Roche catalog no. 09420398001), 200 nM of IDT for Illumina DNA/RNA UD Indexes (Illumina, catalog no. 20026121, 20026930, and 20043019), and 5 µL of template, for a final reaction volume of 50 µL. Duplicate reactions that amplified were pooled together and quantified with Kapa library quantification kit (Kapa Biosystems catalog no. KK4824) before equimolar sample mixing. Libraries were concentrated and cleaned using AMPureXP beads (Beckman Coulter) following each amplification step. The final library was quantified using a High Sensitivity D1000 Tapestation Chip (Agilent catalog nos. 5076-5585, 5067-5584) and Qubit dsDNA Quantification Assay Kit (Thermo Fisher catalog no. Q32851). Sequencing was performed using the Illumina MiSeq platform using V2 chemistry kit (Illumina, PN:MS-103-1003) and 2x300 bp paired-end sequencing. 15% PhiX v3 was added to each sequencing run.

*Fungal 18S and ITS gene amplicon data processing*
Amplicon sequencing data was primarily analyzed using qiime2 (3). Primer sequences and low-quality regions were removed using Cutadapt (4) using linked primers to eliminate read-through of short amplicons. Resulting reads were denoised using DADA2 (5) and taxonomically-classified using a Naive Bayes classifier (sklearn; (6) trained on a database of amplicon sequences obtained by performing in silico PCR (ecoPCR; (7)) on NCBI's RefSeq and GenBank fungal databases (8, 9).

*Illumina Shotgun sequencing*
Extracted DNA was prepared for sequencing as previously described(10) using Illumina DNA prep kit (catalog no. 20018704). Estimations of DNA input were made using fungal 18S primers, the Zymo Femto Human DNA Quantification Kit (catalog no. E2005), and Qubit dsDNA Quantification Assay Kit (Thermo Fisher catalog no. Q32851). For samples with DNA concentrations below Illumina's recommended input, additional PCR cycles were added to the amplification step based on DNA input. Finished libraries were quantified using Qubit dsDNA Quantification Assay Kit (Thermo Fisher catalog no. Q32851) and a High Sensitivity D1000 Tapestation Chip (Agilent catalog nos. 5076-5585, 5067-5584).

*Whole Genome Amplification and PacBio HiFi Sequencing*
Prior to PacBio HiFi sequencing, DNA from selected low-biomass samples was subjected to primary template-directed amplification (PTA) by BioSkryb Genomics (Durham, NC, USA) using the ResolveDNA Long Read workflow. The amplified products were returned to our laboratory and used for downstream PacBio SMRTbell library preparation. Libraries were prepared using the SMRTbell Prep Kit 3.0 (Pacific Biosciences) according to the manufacturer's protocol for metagenomic samples, with minor modifications for low-biomass, pre-amplified DNA. Briefly, the short-read eliminator and DNA-shearing steps were omitted. DNA was first cleaned with 1× SMRTbell cleanup beads and eluted in PacBio Low TE buffer. Libraries then underwent repair and A-tailing, ligation of indexed SMRTbell adapters, post-ligation cleanup, nuclease treatment, and a final 1× SMRTbell cleanup. Bead-based size selection was not performed. Completed libraries were pooled and submitted for quality control and PacBio HiFi sequencing at the University of California, Irvine, on a PacBio Revio instrument. Circular consensus (HiFi) reads were generated and barcode demultiplexing was performed in SMRT Link.

**Figures**

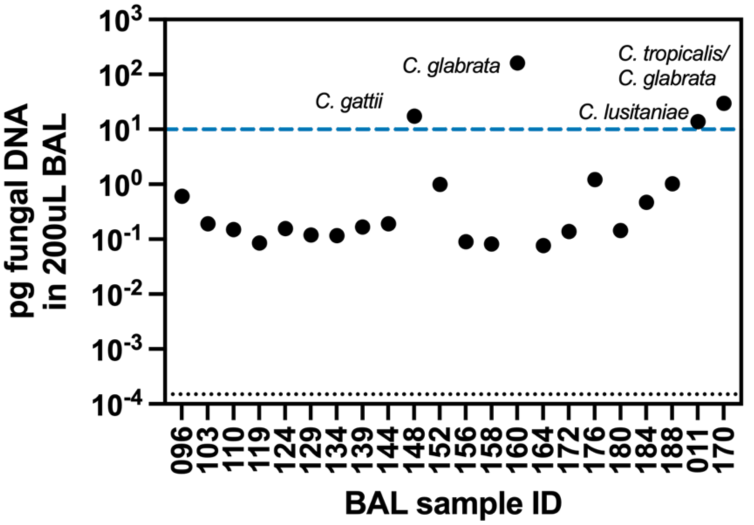


**Figure S1: Summary of BAL samples collected in this study.** The fungal DNA extracted from 200µL of BAL after fMEM is plotted on the y axis for 22 samples collected in this study. The black horizontal dotted line indicates the extraction blank. The blue horizontal dotted line indicates the threshold drawn to proceed with sequencing (>10 pg fungal DNA in 100uL elution from 200uL BAL specimen). The four BAL samples used in this study (011, 148, 160, and 170) are labeled with the dominant fungal species name(s) as determined by culture. Quotations are used around the species name to signify the classification of the isolate given by the clinic.

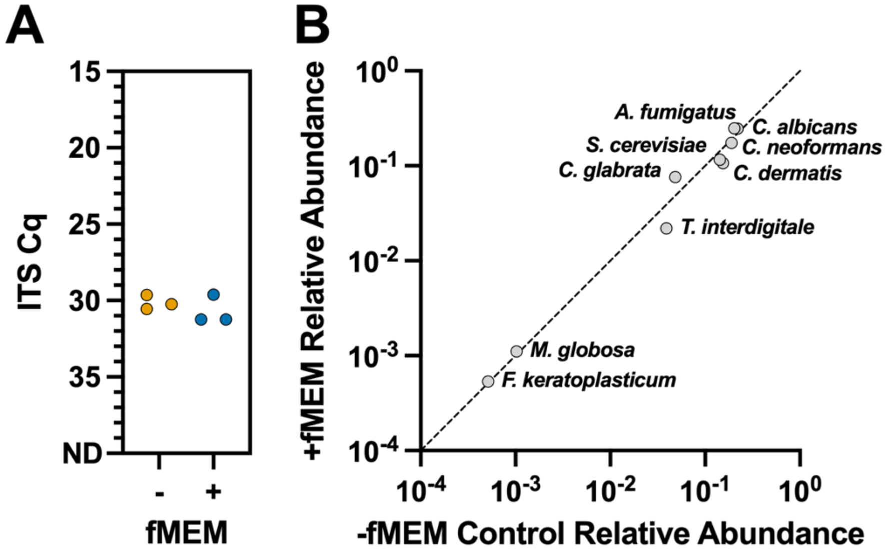


**Figure S2: fMEM preserves fungal community composition as shown by ITS qPCR and amplicon sequencing, similar to the 18S shown in figure 2.** (A) The ITS Cq of the 10-taxa community mix with (blue) and without (orange) fMEM processing. (B) The relative abundance of each fungal taxon (*Aspergillus fumigatus, Cryptococcus neoformans, Trichophyton interdigitale, Penicillium chrysogenum, Fusarium keratoplasticum, Candida albicans, Candida glabrata, Malassezia globosa, Saccharomyces cerevisiae, Cutaneotrichosporon dermatis*) in the 10-taxa community mix after fMEM treatment as a factor of its relative abundance before fMEM treatment, as measured by ITS amplicon sequencing. The relative abundances here reported for ITS are not the same as what is reported for 18S in Fig 2B due to mismatches in fungal primers against specific taxa.

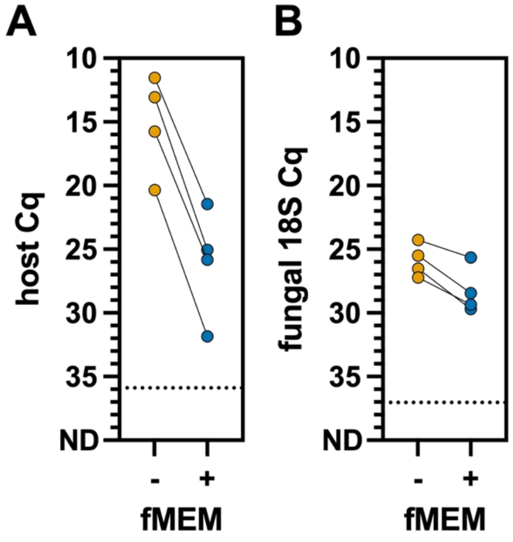


**Figure S3: fMEM reduces human DNA by 1000x while preserving fungal DNA within 10x.** (A) The human DNA load in BAL fluid (measured with Zymo Human Femto qPCR, see methods) for seven samples with (blue) and without (orange) fMEM treatment. (B) The fungal DNA load in BAL fluid (18S Cq) for five fungal culture-positive samples with (blue) and without (orange) fMEM treatment. The horizontal dotted black lines indicate the Cq of the extraction blank negative control.

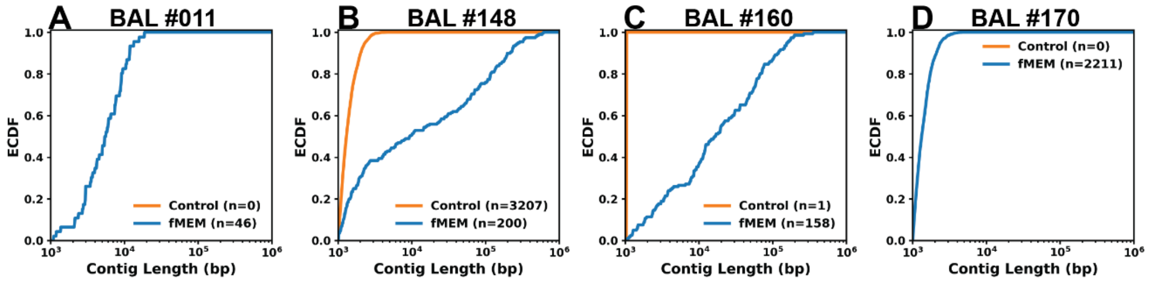


**Figure S4: Fungal contigs assembled from individual BAL samples.** Number of fungal contigs (>1kb) and ECDF of contig length (bp) across four BAL samples with (blue) and without (orange) fMEM processing. (A) BAL #011 *Candida lusitaniae*, (B) BAL #148 *Cryptococcus gattii*, (C) BAL #160 *Candida glabrata*, and (D) BAL #170 *Candida glabrata*/*Candida tropicalis*. Sample #170 contained two fungal isolates.

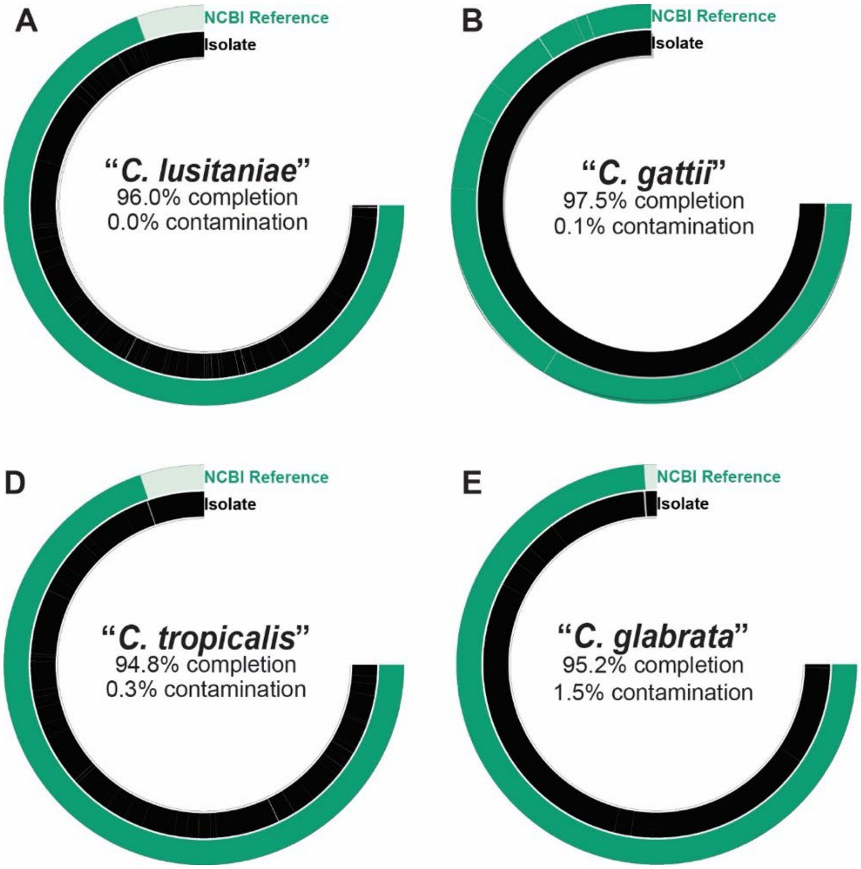


**Figure S5: Fungal genomes assembled from isolates cultured from BAL samples compared to NCBI reference genomes.** For each BAL sample, the fungal genomes assembled from the clinical isolates (black) and scaffolded using the NCBI reference genome (green). (A) "*Candida lusitaniae"* from BAL #011 is being compared to *Clavispora lusitaniae* (GCF_000003835.1), (B) "*Cryptococcus gattii"* from BAL #148 is being compared to *Cryptococcus decagattii* (GCF_036417295.1), (C) "*Candida glabrata"* from BAL #160 is being compared to *Nakaseomyces glabratus* (GCF_000002545.3), (D) "*Candida tropicalis"* from BAL #170 is being compared to *Candida tropicalis* (GCF_000006335.3), and (E) "*C. glabrata"* BAL #170 is being compared to *Nakaseomyces glabratus* (GCF_000002545.3). Percent completion and contamination as determined by BUSCO are stated for each of the isolate genomes. Quotations are used around the species name to signify the classification of the isolate given by the clinic. Sample #170 contained two fungal isolates.

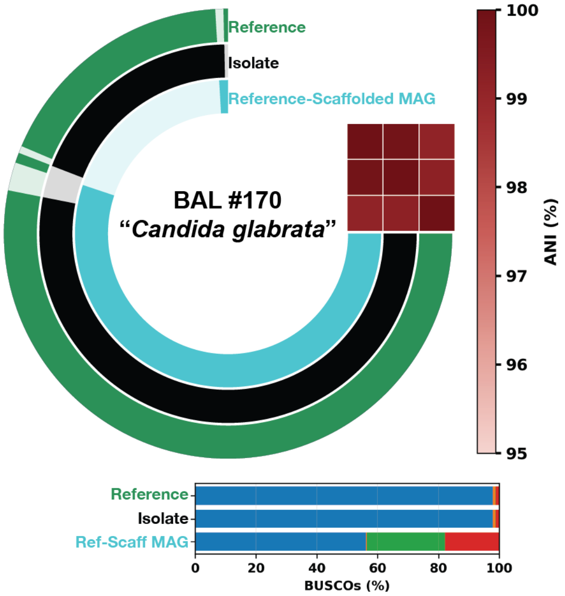


**Figure S6: fMEM processing enables reference-based MAG assembly directly from BAL samples when *de novo* assembly is not possible.** The reference-scaffolded MAGs (light blue) are depicted alongside the assemblies derived from the isolates (black) and the closest-related NCBI reference genome (ATCC 2001; green) for *Candida glabrata* from BAL #170. Percentage completion and percentage contamination, as determined by BUSCO, are stated for each of the reference-scaffolded MAGs. Average nucleotide identity (ANI) between each of the three genomes is shown as a matrix in red. Quotations are used around the species names to signify the classification of the isolate assigned by the clinic.

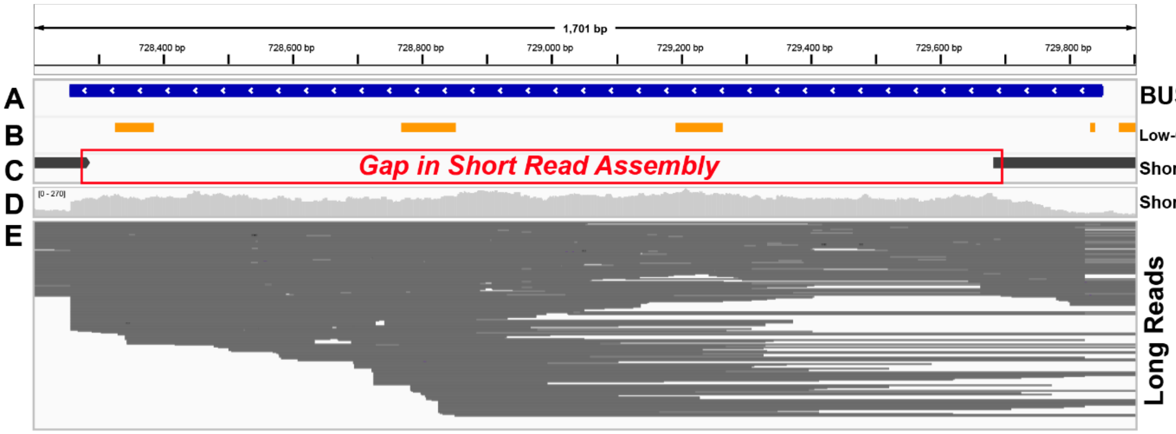


**Figure S7: Long-read sequencing restores a BUSCO gene missed by short-read sequencing.**
The TAF6 BUSCO gene (A) contains low-complexity sequences, detected by DUST (B). The short-read assembly (C) fails to recover this region despite high depth of coverage at the locus (D). Long reads span the low-complexity break (E), enabling the gene to be resolved in the hybrid assembly.

**Table S1: Distribution of Illumina sequencing reads from BAL samples before and after fMEM.**

| sample (isolate) | fMEM | total reads | # human reads | % human reads | # bacterial reads | % bacterial reads | # fungal reads | % fungal reads | # unclassified reads | % unclassified reads |
|---|---|---|---|---|---|---|---|---|---|---|
| **BAL #011 (C. lusitaniae)** | **-** | **40573734** | **39789705** | **98.067644%** | **27740** | **0.068369354%** | **7350** | **0.018115168%** | **748939** | **1.8458715%** |
| **BAL #011 (C. lusitaniae)** | + | **36303991** | **29599962** | **81.533631%** | **5737368** | **15.803684%** | **40075** | **0.11038731%** | **926586** | **2.5522979%** |
| **BAL #148 (C. gattii)** | **-** | **57260744** | **56039859** | **97.867850%** | **19172** | **0.033481926%** | **142911** | **0.24957936%** | **1058802** | **1.8490888%** |
| **BAL #148 (C. gattii)** | + | **8999231** | **3088702** | **34.321844%** | **940446** | **10.450293%** | **4602943** | **51.14818144%** | **367140** | **4.0796819%** |
| **BAL #160 (C. glabrata)** | **-** | **43077723** | **42301858** | **98.198918%** | **6658** | **0.015455784%** | **23909** | **0.055502005%** | **745298** | **1.7301239%** |
| **BAL #160 (C. glabrata)** | + | **50122420** | **44220293** | **88.224577%** | **189221** | **0.377517686%** | **4689936** | **9.3569624%** | **1022970** | **2.0409430%** |
| **BAL #170 (C. tropicalis/C. glabrata)** | **-** | **29619625** | **28940949** | **97.708695%** | **28008** | **0.094558928%** | **7234** | **0.024422997%** | **643434** | **2.1723232%** |
| **BAL #170 (C. tropicalis/C. glabrata)** | + | **102298040** | **76334472** | **74.619682%** | **22470837** | **21.966048%** | **309935** | **0.30297257%** | **3182796** | **3.1112971%** |

**Table S2: Summary statistics for PTA-derived PacBio sequencing libraries.** Values include read counts, read N50, estimated chimera rate, proportion of fungal reads, mean coverage relative to the reference genome, and the fraction of the MAG covered at ≥1X and ≥5X depth.

| Sample | Number of Reads | Read N50 | Percent Chimeras | Percent Fungal Reads | Mean Depth Over Reference | % MAG Covered at 1X | % MAG Covered at 5X |
|---|---|---|---|---|---|---|---|
| **BAL #011** | 774399 | 4688 | 1.71 | 2.90 | 26.8 | 85.1 | 75.0 |
| **BAL #148** | 692457 | 4608 | 0.06 | 70.3 | 113 | 87.4 | 83.3 |
| **BAL #160** | 554255 | 4657 | 0.05 | 21.8 | 42.6 | 96.7 | 95.5 |

**Table S3: BUSCO completeness and gap statistics for (short/hybrid) assemblies.** BUSCO results and gap statistics are shown for assemblies generated using short-read data alone ("Short") or hybrid assemblies incorporating long-read data ("Hybrid"). Because short-read assemblies were already of high quality, BUSCO completeness metrics change only modestly after hybrid assembly. In one case (BAL_160), hybrid assembly slightly increases BUSCO completeness and reduces both fragmented and missing BUSCOs, while gap statistics show small reductions in total gap length.

| Sample | Complete (Short) | Complete (Hybrid) | Fragmented (Short) | Fragmented (Hybrid) | Missing (Short) | Missing (Hybrid) | Total Gaps (bp; Short) | Total Gaps (bp; Hybrid) | % Gaps (Short) | % Gaps (Hybrid) |
|---|---|---|---|---|---|---|---|---|---|---|
| **BAL#011** | 723 | 723 | 6 | 6 | 29 | 29 | 283,102 | 282,240 | 2.35 | 2.34 |
| **BAL#148** | 719 | 719 | 7 | 7 | 32 | 32 | 603,171 | 603,171 | 3.37 | 3.37 |
| **BAL#160** | 720 | 722 | 6 | 5 | 32 | 31 | 395,190 | 392,575 | 3.17 | 3.15 |

**Author Contact Information and ORCiDs:**


| Name | Email | ORCID |
|---|---|---|
| Michael K. Porter | | 0000-0002-0777-7563 |
| Reid T. Akana | | 0000-0003-4422-587X |
| Anna E. Romano | | 0000-0002-7148-0668 |
| Xinyue (Penny) Pei | | 0009-0003-9840-6243 |
| Bishoy Kamel | | 0000-0003-2934-3827 |
| Sajeet F. Haridas | | 0000-0002-0229-0975 |
| Kurt LaButti | | 0000-0002-5838-1972 |
| Igor V. Grigoriev | | 0000-0002-3136-8903 |
| Natalie J. Wu-Woods | | 0000-0001-5070-9091 |
| Omai Garner | | 0000-0002-7366-2692 |
| David Underhill | | 0000-0002-2989-658X |
| Rustem F. Ismagilov | | 0000-0002-3680-4399 |

**Detailed Author Contributions:**
**MKP**: Designed the overall study, performed experiments, analyzed wet lab experimental data, generated Figures 1, 2, S1, S2, and S3, and helped draft the manuscript.
**RTA**: Assisted with overall study design, helped with BAL nucleic acid extractions, performed bioinformatic analyses, generated Figures 3–6, Figures S4–S6, and Tables S1–S3, and drafted the manuscript. Coordinated manuscript review with co-authors and led manuscript revisions.
**AER**: Contributed to the implementation of the fungal amplicon sequencing workflow, performed amplicon sequencing, provided technical guidance for experimental design and data interpretation, assisted with background literature research, helped with amplicon and shotgun library preparation, assisted with overall study design, and edited the manuscript and figures.
**XPP**: Helped with amplicon and shotgun library preparation.
**BK**: Provided guidance on data analysis and interpretation, analyzed sequencing data, assembled genomes and MAGs for Figures 4, 5, 6, S5, and S6, and provided functional annotations for all assembled genomes and MAGs.
**SFH**: Assisted with annotation of assembled genomes and MAGs used in Figures 4–6, S5, and S6.
**KL:** Provided guidance on data interpretation.
**IVG**: Provided guidance on data analysis and interpretation.
**NWW**: Helped with amplicon and shotgun library preparation and assisted with overall study design.
**OG**: Provided guidance on study design and provided BAL samples and cultured isolates for analysis.
**DU**: Provided guidance on experimental design and provided cell cultures for in vitro validation shown in Figure 2.
**RFI**: Served as investigator, provided feedback on study design, provided leadership, technical guidance, and oversight, and obtained funding for the study.
